# Integrating genomic resources to present full gene and promoter capture probe sets for bread wheat

**DOI:** 10.1101/363663

**Authors:** Laura-jayne Gardiner, Thomas Brabbs, Alina Akhunova, Katherine Jordan, Hikmet Budak, Todd Richmond, Sukwinder Singh, Leah Catchpole, Eduard Akhunov, Anthony Hall

## Abstract

**Background:** Whole genome shotgun re-sequencing of wheat is expensive because of its large, repetitive genome. Moreover, sequence data can fail to map uniquely to the reference genome making it difficult to unambiguously assign variation. Re-sequencing using target capture enables sequencing of large numbers of individuals at high coverage to reliably identify variants associated with important agronomic traits.

**Results:** We present and validate two gold standard capture probe sets for hexaploid bread wheat, a gene and a promoter capture, which are designed using recently developed genome sequence and annotation resources. The captures can be combined or used independently. We demonstrate that the capture probe sets effectively enrich the high confidence genes and promoters that were identified in the genome alongside a large proportion of the low confidence genes and promoters. Finally, we demonstrate successful sample multiplexing that allows generation of adequate sequence coverage for SNP calling while significantly reducing cost per sample for gene and promoter capture.

**Conclusions:** We show that a capture design employing an ‘island strategy’ can enable analysis of the large gene/promoter space of wheat with only 2×160 Mb probe sets. Furthermore, these assays extend the regions of the wheat genome that are amenable to analyses beyond its exome, providing tools for detailed characterization of these regulatory regions in large populations.

## Background

The allohexaploid (AABBDD) wheat genome is 17Gb in size and derived from three diploid progenitor genomes. The AA genome is from *Triticum urartu*, the BB is likely to be of the Sitopsis section (includes *Aegilops speltoides*), and the DD from *Aegilops tauschii* (Brenchley *et al.*, 2012). AABB tetraploids appeared less than 0.5 million years ago after an initial hybridization event (Dvorak *et al.*, 2006). It is thought that Emmer tetraploid wheat developed from the domestication of such natural tetraploid populations. The hexaploid wheat that we have today formed around 8000 years ago by the hybridization of the unrelated diploid wild grass *Aegilops tauschii* (DD genome) with the tetraploid *Triticum turgidum* or Emmer wheat (AABB genome) (Dubcovsky and Dvorak, 2007).

It is expensive to perform whole genome sequencing in wheat because of its vast genome, polyploid nature and high repetitive content. To reduce this complexity and to make re-sequencing of wheat more cost effective, we can utilize approaches such as: Restriction site Associated sequencing or RAD-seq (Baird *et al.*, 2008), transcriptome sequencing (De Wit *et al.*, 2015) and sequence capture. Sequence capture typically combines probe hybridization to capture specific genome sequences in solution with sequencing of the captured fragments. The ability to design and implement specifically targeted probe sets has clear advantages for the analysis of variation across the genome. With annotated high-quality wheat genome sequences now available, it has become possible to design such capture probe sets for wheat and to use them to accurately analyze the genome (Clavijo *et al.*, 2017; IWGSC). Sequence capture combines genotyping with de novo SNP/CNV discovery to allow allele mining and identification of rare variants. To date, such diversity has been profiled in wheat using capture probe sets that have not been able to make use of the recent advances in wheat genome sequencing and annotation. Most of the diversity studies have implemented either cDNA/exon-based probe sets of 56 and 84 Mb (Winfield *et al.*, 2012; Krasileva *et al.*, 2016) or the gene-based probe set of 107 Mb (Jordan *et al.*, 2015; Gardiner *et al.*, 2016). Aligning the 107 Mb capture probe set to the current wheat genome annotations (BLASTN, e-value 1e-05, identity 95%, minimum length 40bp), we can see that it represents only 32.9% of the high confidence gene-set or 21.2% of the gene-set plus promoters defined as 2Kbp upstream (Clavijo *et al.*, 2017). Similarly aligning the 84 Mb capture probe set to the current wheat genome annotations, we can see that it represents only 32.6% of the high confidence gene-set or 20.4% of the gene-set plus promoters. Promoter and intron sequence has previously been largely missing from capture probe sets alongside the newly characterized genes that the recent improved reference sequences have defined. There is therefore a need for an updated “gold-standard” gene capture probe set for wheat, based on the current high confidence gene-models, that can be adopted by the community.

Here, we present a gene capture probe set, which was created by integrating the current annotated wheat genome reference sequences to define a comprehensive “gold-standard” gene design space for wheat. We use an island strategy, carefully spacing probes with, on average, 120 bp gaps across the design space to maximize sequencing coverage of our targets. We have also developed a comprehensive promoter capture probe set for wheat that takes 2 Kb upstream of the annotated genes and will facilitate global investigation to fully characterize these regulatory regions. Since approximately half of the genetic variation that associates with phenotypic diversity in maize is found in promoter regulatory regions (Li, X. *et al.*, 2012), it is reasonable to expect a similar scenario for wheat promoter regions that are poorly defined on a global scale; these are regions that need to be explored and more precisely defined across the wheat genome. The gene and promoter captures can be combined or used independently.

In summary: we describe two new wheat NimbleGen SeqCap EZ probe sets (Roche NimbleGen Inc., WI, USA), the first that is tiled across the genic regions of the hexaploid bread wheat genome and the second that is tiled across the promoter regions; we integrate diverse wheat material into the design to allow broad applicability of the probe sets; we validate the capture probe sets using the reference variety Chinese Spring; and we demonstrate the probe sets application to diverse wheat accessions by enriching eight wheat accessions that were generated by International Maize and Wheat Improvement Center (CIMMYT), Mexico. Finally, multiplexing samples into a single capture before sequencing, using barcodes to identify individual samples in the pool, can further reduce costs; we demonstrate successful multiplexing of over 20 samples in a single capture, where we can generate adequate coverage per sample for SNP calling. Our capture probe set designs are publicly available and can also be ordered directly from NimbleGen via the Roche website (http://sequencing.roche.com/en/products-solutions/by-category/target-enrichment/shareddesigns.html).

## Analyses

### Targets of the capture probe sets (Figure 1)

**Figure 1.**
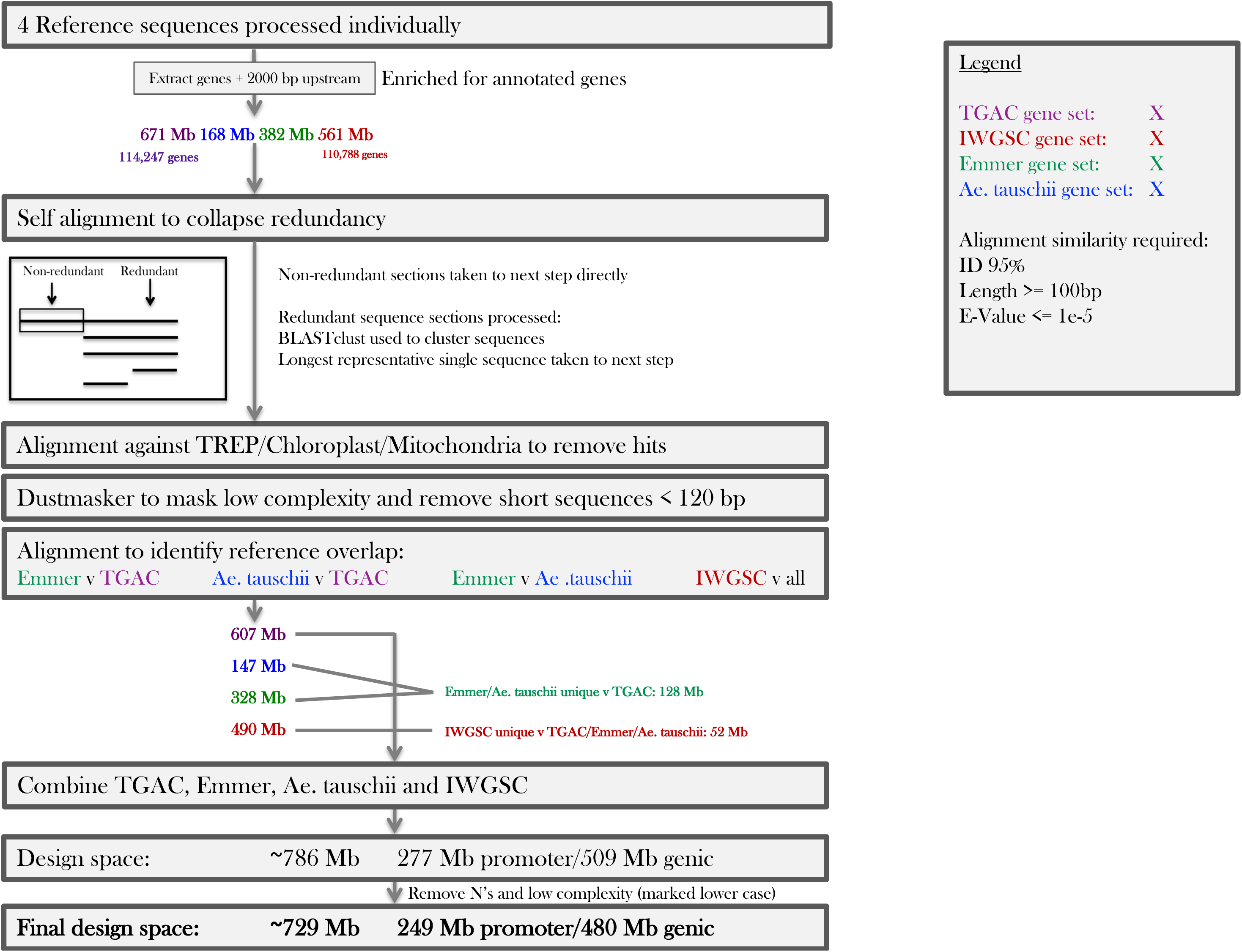
Design of the wheat gene and promoter capture probe sets. Processing of the TGAC Chinese Spring, IWGSC Chinese Spring, Emmer and Ae. tauschii reference sets of gene/promoter sequences to generate a final design space for the wheat gold standard promoter/gene capture probe set that is; non-redundant and high complexity (Methods).

The capture probe sets target high confidence genes and their associated promoters from: the Chinese Spring reference genome, Chinese Spring’s D-genome progenitor *Aegilops tauschii* and its AB-genome progenitor *Triticum turgidum* or Emmer wheat. For Chinese Spring-derived genes two genome sequence annotations were utilized; high confidence genes from The Genome Analysis Centre (TGAC)/Earlham Institute W2RAP pipeline derived reference sequence (Clavijo *et al.*, 2017) and high confidence genes from the International Wheat Genome Sequencing Consortium (IWGSC) RefSeq.v1 genome assembly (unpublished). For *Aegilops tauschii* genes the Luo *et al.* reference sequence was used with a high confidence annotated gene set and for Emmer wheat, high confidence genes from the Avni *et al.* reference sequence were used.

The target space for the gene capture amounted to 447,729,570 bp of target sequence across 114,247 genes and 339,580,651 bp across 110,788 genes for the TGAC and IWGSC references respectively. *Ae. tauschii* derived genes totalled 111,466,178 bp of target sequence across 28,843 genes and for Emmer wheat 252,137,485 bp across 65,005 genes. Only high confidence genes were selected and gene sequence was defined from the beginning of the 5’untranslated region (UTR) to the end of the 3’UTR sequence.

The target space for the promoter capture amounted to 223,409,786 bp of target sequence across 112,999 gene promoters and 221,681,783 bp across 110,788 gene promoters for the TGAC and IWGSC references respectively. *Ae. tauschii* derived promoters totalled 57,177,213 bp of target sequence associated with 28,843 genes and for Emmer 130,075,005 bp associated with 65,005 genes. Promoter sequence was defined as 2000 bp upstream of the transcription start site (TSS). The 2 Kb distance was based on the median distance between the TSS and the first transposon, 1.52 Kb to allow a high likelihood of full promoter sequence capture (Wicker *et al.*, 2018).

In total, the gene and promoter capture design spaces amounted to: 671,139,356 bp for the Chinese Spring TGAC reference, 561,262,434 bp for the IWGSC Chinese Spring reference, 168,643,391 bp for *Ae. tauschii* and 382,212,490 bp for Emmer. The processing of these raw gene/promoter sequences to remove redundancy, repetitive/low-complexity sequence and chloroplast/mitochondrial sequence (Figure 1; Methods) resulted in probe set design spaces of: 606,847,164 bp (TGAC), 490,375,105 bp (IWGSC), 328,407,758 bp (Emmer) and 146,980,738 bp (*Ae. tauschii*). Unique Emmer/*Ae. Tauschii* sequences amounted to 127,651,054 bp and were combined with the TGAC wheat design space. Finally, the unique Chinese Spring IWGSC design space of 51,758,271 bp was also included. This ensured that gene annotation differences between the two main wheat references sequences were accounted for in the capture design space. As anticipated, overlap between the two Chinese Spring reference annotations was high.

The final TGAC/Emmer/Tauschii/IWGSC gene and promoter design space was 785,914,746 bp, of which, 508,889,665 bp was gene and 277,025,081 bp was promoter sequence. The promoter design space included additional micro RNA (miRNA) sequence totalling 953 sequences (208,968 bp). N’s and low complexity space encompassed 56,648,010 bp of the final design space, although this sequence was included for probe design and later used to enable ranking of more or less preferential probes.

### Characteristics of the gene and promoter capture kits

The final gene/promoter design space of 785,914,746 bp was used for probe design (Methods). Typically probes overlap one another to most optimally cover the target design space, however from previous analyses we observed that a single 120 bp probe can enrich up to 500 bp with adequate sequencing coverage (Gardiner *et al.*, 2015). As such, we tiled probes (average size 75 bp), across our design space using an “island strategy” i.e. at intervals of on average 120 bp, to most evenly cover the design-space. The gene and promoter probe set’s predicted performance metrics and designs are summarized in Table 1. Probes in solution bind to their complementary sequence within a DNA library fragment that has typically been sheared to 200-300bp, therefore we estimated design space coverage based on the probe set using shearing sizes for library construction of 200 bp. We anticipate upwards of 90% coverage of both the promoter and gene capture design-spaces with these capture probe sets. Additionally, we visualized the predicted coverage of the Chinese Spring high confidence genes/promoters by their corresponding design spaces; Supplementary Figure S1a and S1b highlight that the captures are likely to provide a comprehensive coverage of their respective targets. It is also evident that the collapse of the gene design space has been more widespread, with many regions of the capture design aligning closely to more than one target region. This is less common for the promoter capture design sequences that are more likely to align to a single target promoter with a longer alignment. This could be indicative of genes being more likely to have shared homology between the sub-genomes of wheat or within gene families compared to promoters that may be more divergent.

**Table 1.**
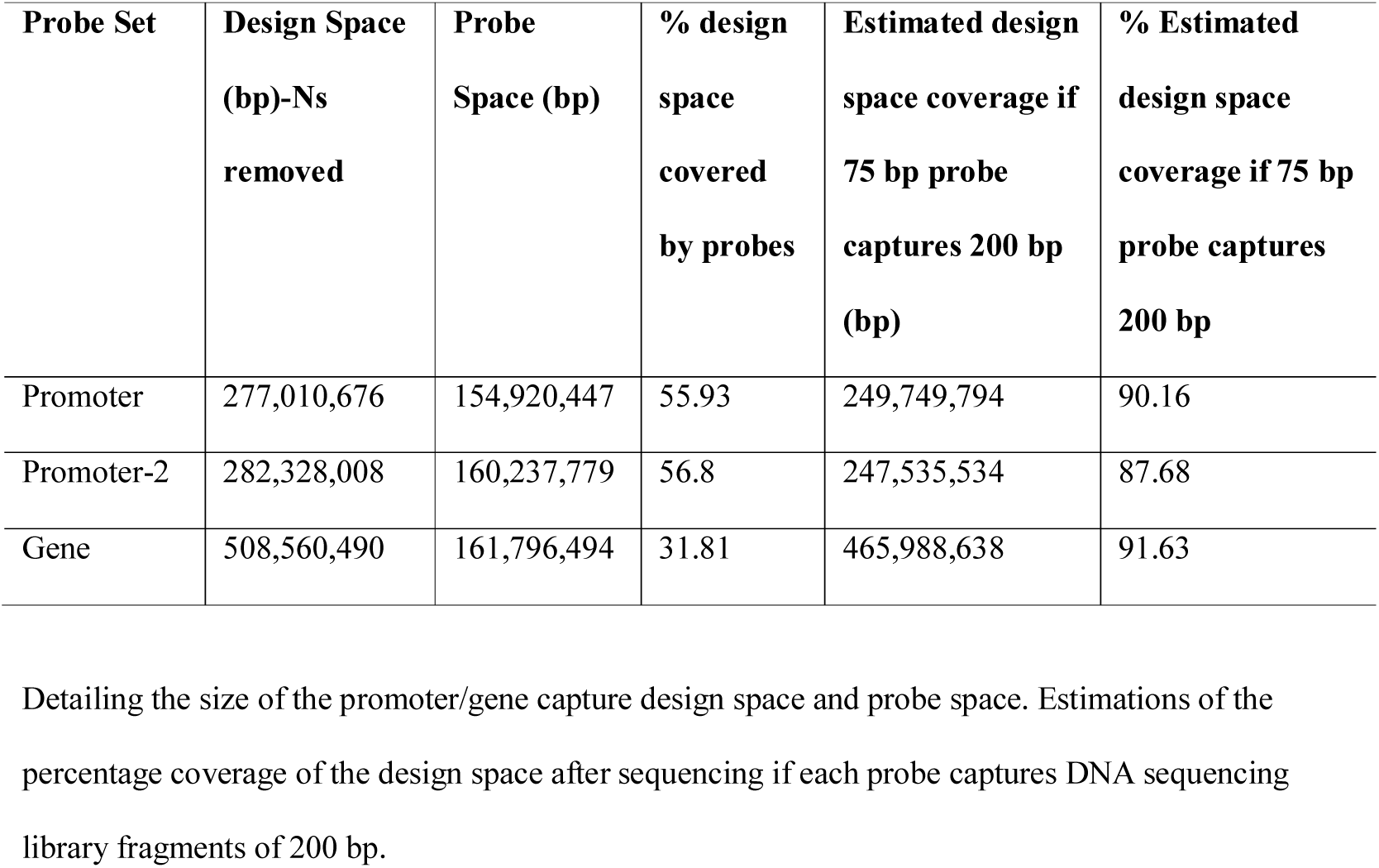
Probe set designs and predicted performance metrics.

| Probe Set | Design Space<br>(bp)-Ns<br>removed | Probe<br>Space (bp) | % design<br>space<br>covered<br>by probes | Estimated design<br>space coverage if<br>75 bp probe<br>captures 200 bp<br>(bp) | % Estimated<br>design space<br>coverage if 75 bp<br>probe captures<br>200 bp |
| --- | --- | --- | --- | --- | --- |
| Promoter | 277,010,676 | 154,920,447 | 55.93 | 249,749,794 | 90.16 |
| Promoter-2 | 282,328,008 | 160,237,779 | 56.8 | 247,535,534 | 87.68 |
| Gene | 508,560,490 | 161,796,494 | 31.81 | 465,988,638 | 91.63 |

Detailing the size of the promoter/gene capture design space and probe space. Estimations of the percentage coverage of the design space after sequencing if each probe captures DNA sequencing library fragments of 200 bp.

### Sequencing coverage after capture of Chinese Spring

We firstly examined capture efficiency using the reference variety of wheat, Chinese Spring, which the majority of the capture design space was based on. We performed promoter and gene captures separately using Chinese Spring DNA from 21-day seedling leaf tissue and sequenced on the HiSeq4000 (Methods). Sequencing data was then aligned and coverage was assessed. It is clear from Table 2 that, irrespective of the reference genome implemented, the majority of reads can be aligned uniquely (average 77.9%) with a low duplicate rate observed (average 4.20%).

**Table 2.**
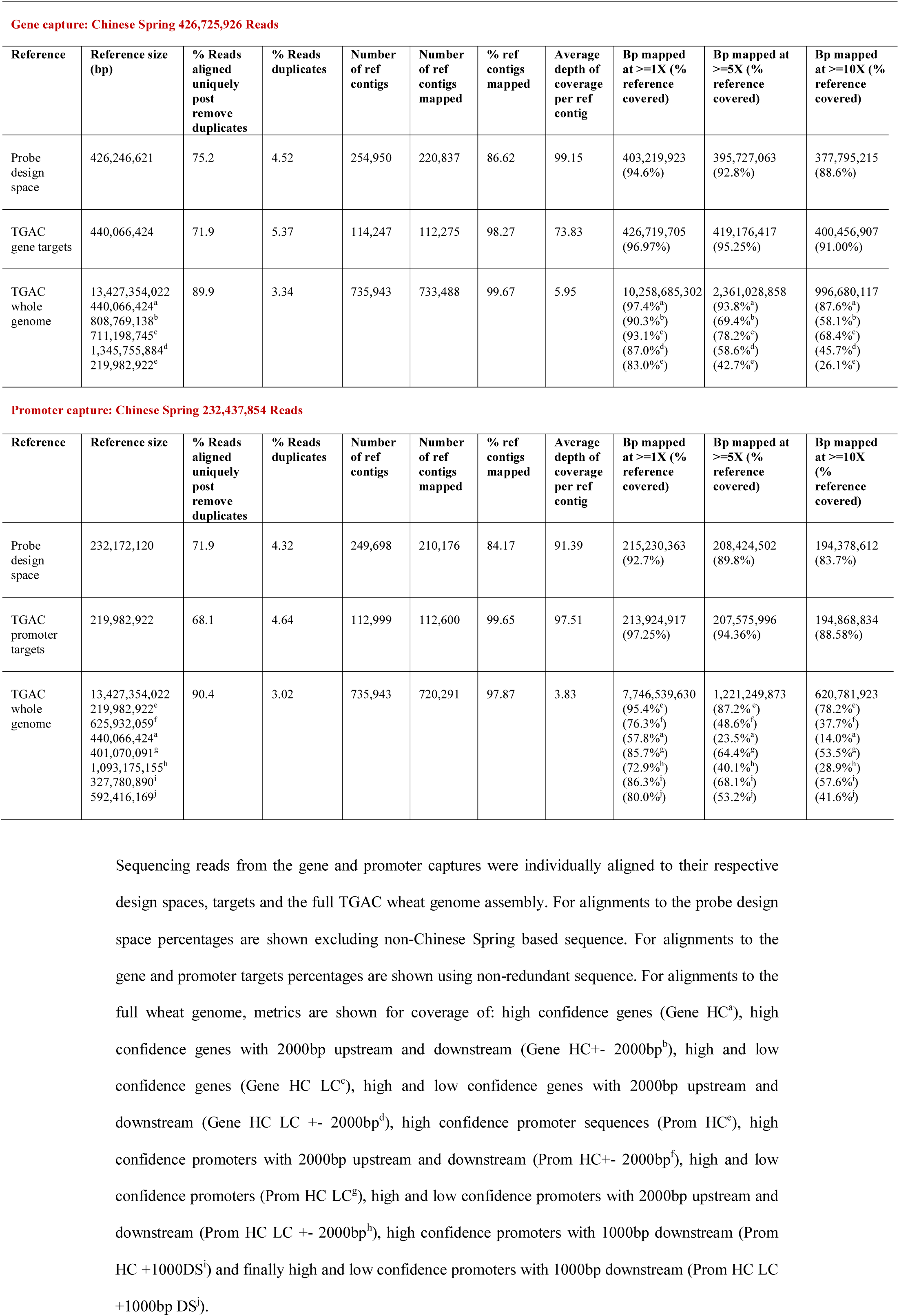
Coverage statistics for Chinese Spring.

Firstly, we aligned promoter and gene captured reads to their respective probe design spaces to determine enrichment efficiency in general i.e. how much of the sequencing data was likely to have been pulled down by the probes (Table 2). 75.2% and 71.9% of reads align to the gene and promoter probe design spaces respectively indicating high on-target enrichment efficiency. For the gene capture probe design space, we saw 94.6% and 92.8% of the design space with coverage at 1X and 5X or more respectively (excluding non-Chinese Spring design space from calculations). Similarly, for the promoter capture design space we saw 92.7% and 89.8% of the design space with coverage at 1X and 5X or more respectively. The performance of the promoter and gene capture platforms exceed our predictions of coverage of 90.2% and 91.6%. Coverage statistics for the probe design spaces were used to identify regions with excessively high coverage defined as more than 10X the average maximum depth of coverage for a region. Only 0.17% of gene and 0.22% of promoter design space regions showed such high coverage and will be removed from subsequent versions of the capture probe sets.

Secondly, we focused on alignments to the full high confidence gene and promoter spaces of Chinese Spring, i.e. only our intended targets, to determine the efficacy of our island approach and design space collapse (Table 2 and Figure 2). For the gene capture we observed a highly comprehensive coverage of 96.97% and 95.25% at 1X and 5X or more respectively. Similarly, for the promoter capture we observed 97.25% and 94.36% coverage at 1X and 5X or more. This demonstrates exceptional performance of the island approach and Figure 2 highlights this ability of the short probes to generate comprehensive coverage using the island approach. We noted that coverage of the full gene and promoter sets actually exceeds that of the probe design space. This is likely due to the full gene and promoter space having a smaller number of contigs (up to 114,247) that are generally longer and encompass a larger base space compared to the probe design space, which has a larger number of contigs (up to 220,837) that are shorter in length and therefore likely to hinder successful mapping of properly paired reads.

**Figure 2.**
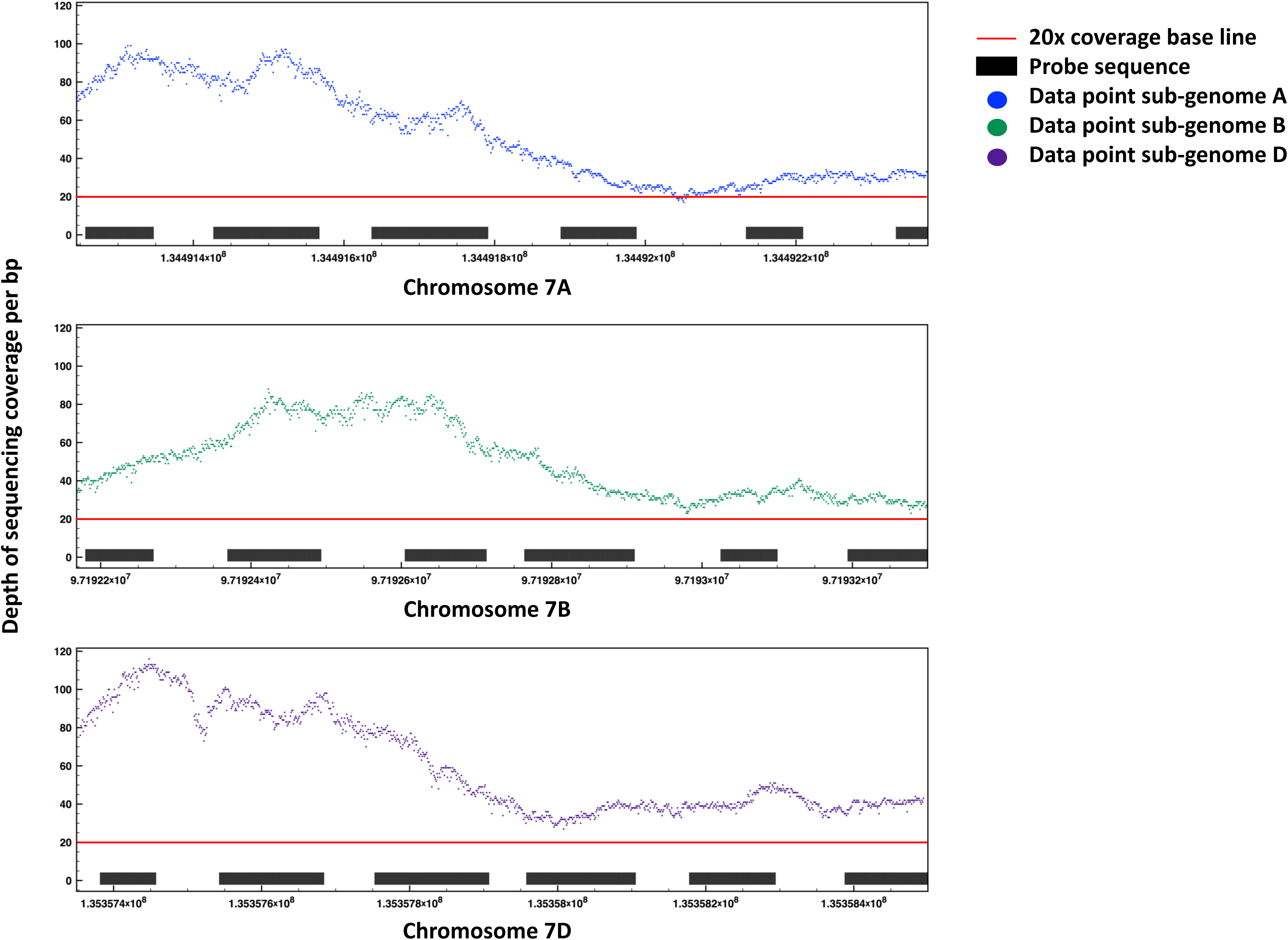
Highlighting coverage of the MYB transcription factor gene triplet using an island probe design approach. The depth of sequencing coverage is shown per base pair across three chromosomal intervals corresponding to a trio of homoeologous genes for the Myb transcription factor (TraesCS7A01G179900 on chr7A at 134491245-134492378bp, TraesCS7B01G085100 on chr7B at 97192168-97193300bp and TraesCS7D01G181400 on chr7D at 135357355-135358494bp).

Finally, to assess off-target sequencing carryover and to ensure unbiased sequencing alignment, we looked at read alignments to the full Chinese Spring genome (Table 2). We observe coverage across 97.4% and 93.8% of the high confidence genic regions at 1X and 5X or more respectively. Furthermore, we see 93.1% and 78.2% coverage at 1X and 5X or more across all high confidence and low confidence genes, resulting in a truly comprehensive gene capture. 113,884 of the high confidence genes (99.7%) showed sequencing coverage, with each gene covered to an average of 97.5% at 1X and 94.5% at 5X or more. The promoter capture performed comparably to the gene capture showing coverage across 95.4% and 87.2% of high confidence promoters at 1X and 5X or more and also coverage of 85.7% at 1X and 64.4% at 5X or more across promoters associated with both high and low confidence genes. Here, a slightly lower coverage of low confidence promoter sequences was observed than for genes, potentially due to more divergent or repetitive promoter associated sequences. 112,824 of high confidence promoters (99.8%) showed sequencing coverage, with each promoter covered to an average of 93.6% at 1X and 85.5% at 5X or more. Within the design space of the promoter capture we included miRNA sequence totalling 953 sequences (208,968 bp). We observed coverage across 92.72% of these sequences with an average depth of 34.99X with as little as 47 million sequencing paired-end reads (23.5 million read clusters).

Using the information from the Chinese Spring sequencing validation of the gene capture probe set, we were able to develop an extended version of the promoter capture probe set that includes 5’UTR sequence (Promoter-2). The 5’UTRs for which we gained coverage of >10X across >99% of the 5’UTR sequence were identified and up to 2 probes per 5’UTR were added to the promoter capture. This resulted in the addition of 5’UTR probes that were associated with 49,034 high confidence genes. This provides an enhanced promoter capture probe set that overlaps the first probe set with the addition of the 5’UTR.

To assess the compatibility of our capture probe set with different Chinese Spring reference genome sequences, we performed further read alignments to the full IWGSC Chinese Spring genome (RefSeqV1, Supplementary Table S1). 109,862 of high confidence genes (99.2%) showed sequencing coverage, with each gene covered to an average of 98.0% at 1X and 94.2% at 5X or more. In addition, 109,986 of high confidence promoters (99.3%) showed sequencing coverage, with each promoter covered to an average of 90.4% at 1X and 79.0% at 5X or more respectively. These statistics are highly comparable to the outcome using the TGAC reference and highlight the large degree of overlap that is seen between the TGAC and IWGSC Chinese Spring reference gene sets, aside from small regions of inverted duplications (Supplementary Figure S1c). Since we do not see a significant difference in coverage between the TGAC and IWGSC reference sequences this confirms our ability to capture much of the regions differing between the two references.

Using the IWGSC Chinese Spring genome that is ordered into chromosome pseudomolecules we can visualize the genome-wide average coverage of genes and promoters (Supplementary Figure S2). We see no notable bias in coverage depth or distribution between the sub-genomes of wheat or otherwise. Coverage is consistent across the vast majority of the gene and promoter space with baseline averages of 34.7X and 21.0X coverage. For the gene capture, the coverage coefficient of variation (CV) is 0.87, while for the promoter capture it is 0.79; distributions with CV < 1 are considered low-variance and as such coverage is largely uniform across the respective target spaces.

### Capturing and sequencing regions not included in the target space

Overall both captures perform well, we can typically gain >5X coverage across >90% of their intended targets and on average >20X coverage. It was noted across both captures that there was a significant proportion of low level coverage that fell outside of high and low confidence genes, promoters and sequences in their immediate vicinities (+/- 2000bp). This is visible in the ~20% difference in reads aligning uniquely to the whole wheat genome but not to the TGAC gene/promoter targets. This sequence is thought to be non-enriched carryover contamination and as such could be limited with increased washes during the capture protocol. There is also the possibility that this may be a result of our high level “over-sequencing” of the libraries here and as such we hypothesize that the off-target sequence will become less prominent at lower sequencing depths; we already see an increase in on-target sequence of 1.1% as we decrease read coverage from 440 to 100 million sequencing reads for the gene capture.

### Determination of minimum sequencing requirements

The Chinese Spring data that we have used to validate the capture probe sets originated from a single gene and a single promoter capture assay, however, each capture combined four barcoded technical replicate Chinese Spring libraries. Using different combinations of these four replicate libraries (all four, three, two or one) we were able to bioinformatically reduce the number of sequencing reads in our analyses to determine the minimum sequencing requirements for coverage of the targets. Looking at the coverage of target regions with varying sequencing read numbers (Supplementary Table S2 and Supplementary Figure S3), it is evident that increasing the number of sequencing reads increases coverage of target regions. However, there are clear saturation points for each capture probe set where further sequencing input has little to no effect on increasing target coverage. These saturation points guide our recommended sequencing levels for optimal return on investment and comprehensive coverage of capture targets at a minimum of 5X, which is desirable for SNP calling: 200-300 million paired-end reads (100-150 million read clusters) for gene capture and 150-200 million paired-end reads for promoter capture (75-100 million read clusters). In Table 3 we have outlined a sliding scale of sequencing levels alongside the varying depths of coverage that they generate for the target sequences to guide user requirements (Table 3).

**Table 3.** Sequencing recommendations for gene and promoter capture probe sets.

| **Capture**<br>**Probe set** | **Approximate**<br>**Read**<br>**Number**<br>**Required**<br>**with**<br>**standard**<br>**protocol** | **Approximate**<br>**Read**<br>**Number**<br>**Required**<br>**with**<br>**optimized**<br>**protocol** | **Expected**<br>**%**<br>**coverage**<br>**of target**<br>**(>=1X)** | **Expected**<br>**%**<br>**coverage**<br>**of target**<br>**(>=5X)** | **Expected**<br>**%**<br>**coverage of**<br>**target**<br>**(>=10X)** | **Average**<br>**coverage**<br>**across**<br>**target**<br>**region\*** |
| --- | --- | --- | --- | --- | --- | --- |
| Gene | 100,000,000 | 55,000,000 | 94.3 | 69.8 | 35.4 | 9.05 |
| Gene | 200,000,000 | 105,000,000 | 96.4 | 86.9 | 68.3 | 17.13 |
| Gene | 300,000,000 | 160,000,000 | 97.1 | 91.8 | 81.6 | 25.42 |
| Gene | 400,000,000 | 210,000,000 | 97.4 | 93.8 | 87.6 | 34.06 |
| Promoter | 50,000,000 | 30,000,000 | 87.3 | 43.4 | 9.9 | 5.27 |
| Promoter | 100,000,000 | 55,000,000 | 93.2 | 78.2 | 52.6 | 12.05 |
| Promoter | 150,000,000 | 80,000,000 | 94.2 | 83.3 | 66.3 | 15.82 |
| Promoter | 200,000,000 | 105,000,000 | 95.4 | 87.2 | 78.2 | 21.59 |

Projected coverage of gene and promoter capture target sequence (high confidence gene and promoter sequences respectively) with varying numbers of sequencing reads. Also shown is the predicted read number requirements to achieve the same coverage using our optimized capture protocol (numbers rounded to the nearest 5 million reads). Read numbers are for total number of paired-end reads and
should be halved to get the number of read clusters. \*Target region is defined as all gene or promoter sequences that the probe sets are tiled across i.e. including padding between probes.

### Mulitplexing and sequencing to generate comprehensive coverage

Multiplexing DNA from multiple wheat lines and enriching them in a single capture reaction before sequencing can decrease costs. Using barcodes to label individual samples in the multiplexed pool we are able to distinguish individual samples post-sequencing and quantify the efficacy of the enrichment.

Firstly, we multiplexed eight different samples per gene and promoter capture to compare performance metrics with our previous analysis using a single sample per capture. For this analysis we used eight wheat accessions that were generated by CIMMYT, Mexico and sequenced the gene capture multiplexed pool to a depth of 800 million paired-end reads (~ 100 million paired-end reads per sample or 50 million read clusters) and the promoter capture pool to 600 million paired-end reads (~ 75 million reads per sample or 37.5 million read clusters). We performed read alignments for the eight samples to the full Chinese Spring genome (Supplementary Table S3 and S4). Uniform and successful enrichment of the eight samples was observed with both the gene and promoter captures. All samples show a high percentage of reads aligned on target (77.25% and 58.47% on average for the gene and promoter captures respectively) with low variation between samples represented by interquartile ranges of less than 5% (Figure 3). CVs for the gene and promoter capture were 0.68 and 0.59 respectively; these values are considered low-variance and as such coverage is largely uniform across the respective target spaces. All samples covered the gene target regions at a minimum of 5X to between 69.2-73.1% and the promoter target regions at a minimum of 5X to between 62.7-70.4% (Supplementary Tables S3 and S4). For each of the samples, coverage of target regions was higher than the expected coverage that was predicted based on the depth of sequencing from the Chinese Spring enrichment (Supplementary Figure S4).

**Figure 3.**
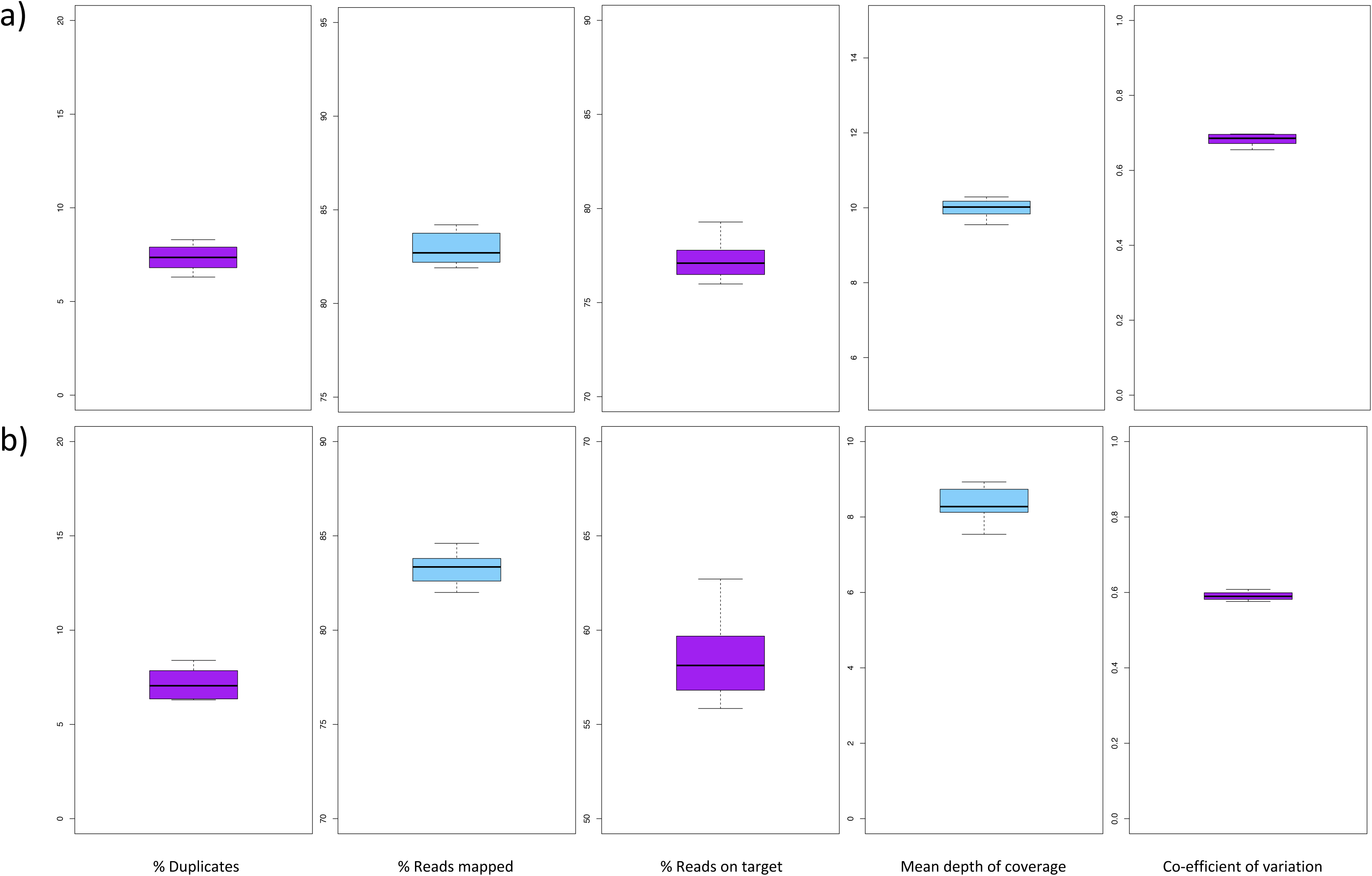
Summary statistics for the 8-plex gene and promoter capture tests. We performed read alignments for the eight CIMMYT samples to the full Chinese Spring genome. For the **(a)** gene capture and **(b)** promoter capture probe sets, from left to right, we show box and whisker plots for: the percentage of sequencing reads per sample that were identified as duplicates, the percentage of reads mapping uniquely to the whole genome reference sequence, the percentage of reads defined as ‘on target’ i.e. align to the capture probe design space, the mean depth of coverage per sample and the coefficient of variation per sample.

Secondly, we validated our promoter-2 capture probe set that includes 5’UTR sequence whilst also multiplexing a larger number of samples for capture (22 samples). For this analysis we used a diverse set of wheat lines that were selected based on genotyping with the 9K iSelect array (Cavanagh *et al.*, 2013). We sequenced the promoter-2 capture multiplexed pool to an average depth of ~46 million paired-end reads per sample (23 million read clusters) and aligned on average 85.5% of reads uniquely to the reference genome. Across a representative subset of the samples, the average depth of coverage for the promoter high confidence target regions ranged from 6.2-6.9X with 13.5-17.1% of this space covered at a minimum of 10X. These metrics surpass our expected depth of coverage on target using 50 million paired-end reads where we predicted an average coverage of 5.3X and 9.9% coverage at a minimum of 10X (Table 3). We also noted low variation between samples represented by interquartile ranges of less than 5% for coverage of the targets at a minimum of 1X, 5X and 10X. This analysis demonstrates uniform successful enrichment of the samples and that multiplexing more than 20 samples for capture has no detrimental effect.

### Genotyping sensitivity of the capture probe sets

We focused on our 8-plex test, where we sequenced the samples to our recommended sequencing depth for SNP calling, and we identified homozygous SNPs in each of the samples at positions where we saw a minimum of 5X coverage (Methods). On average samples had 1,031,677 SNPs each from the gene capture and 968,640 SNPs each from the promoter capture. Furthermore, when we focus on locations where each of the eight samples either had a SNP identified or else had a minimum of 5X coverage with no SNP, i.e. the reference allele, this resulted in 1,019,556 positions that were available for comparison across the sample set for the gene capture and 869,954 for the promoter capture. This highlights our ability for *denovo* SNP discovery with captured sequencing data, the high level of diversity in the eight CIMMYT lines compared to the Chinese Spring reference and the successful uniform enrichment of these samples despite this diversity.

### Optimizing the capture protocol

Due to the large size of our capture probe sets we performed further optimization of the standard NimbleGen capture protocol to focus our sequencing reads on target as much as possible. We again used Chinese Spring for this analysis in a repeat of our initial quality control of the capture. Here, we noted a reduced fragment size of the sheared DNA to focus the fragments onto the probe design islands and to maximize the coverage of the DNA fragments by sequencing reads (2×150bp reads). We combined both the promoter and gene capture probe sets for analysis and we also implemented an additional wash to remove any remnant off-target fragments (Methods). For this analysis we noted that 57% of the mapped reads were on target i.e. aligned directly to the probe design. This is in line with what we observed previously, with a range of 58.47-77.25% observed across the gene and promoter captures. However, we noted that here, rather than enriching high and low confidence genes there was a bias specifically towards high confidence genes with a 1.9-fold increase the sequence space aligning to these genes compared to previous analyses. This allowed us to lower our original predictions of sequencing requirements for adequate coverage of the high confidence gene set (Table 3).

## Discussion

Sequence capture is rapidly becoming one of the main techniques employed by the wheat research community for re-sequencing of the large complex wheat genome at reduced cost. It allows the identification of previously uncharacterized genetic variation in the form of SNPs and indels in key regions of interest that are typically gene-associated. To date, many studies have implemented either exon or cDNA-based capture probes sets that have not been able to make use of the recent advances in wheat genome sequencing and annotation. Furthermore, promoter and intronic sequence has largely been missing from capture probe sets. Here, we present and validate a gene capture probe set, created by integrating the current annotated wheat genome reference sequences to define a comprehensive “gold-standard” gene design space for bread wheat. We have also developed a comprehensive promoter capture probe set for wheat that covers 2 Kb upstream of the annotated genes and will facilitate global investigation to fully characterize these regulatory regions. An updated version of the promoter capture probe set also includes gene 5’UTRs and so will capture regulatory elements within these regions.

We have demonstrated the use of the capture probe sets to analyze a diverse set of material including pure breeding lines that were generated by CIMMYT, Mexico. We studied the consistency of our data by correlating the sequence coverage depths between independent captures for multiple DNA samples. In addition, we successfully multiplex over twenty samples in a single capture with no drop out of capture efficiency despite the large size of our capture. From multiplexed captures, we can generate adequate coverage per sample for SNP calling resulting in a lower cost per sample for gene and promoter captures. This brings down the cost of re-sequencing the entirety of wheat’s gene associated space. Furthermore, it is likely that, since no reduced capture efficiency was observed with a 20-plex capture, more samples could be multiplexed without a detrimental effect. We have focused on generating a depth of coverage that is adequate for SNP calling, but the potential is there for skim sequencing samples. Skim sequencing generates low coverage for a larger number of lines to allow allele mining at reduced cost and this can be achieved using multiplexing or bulk segregant analysis that we have previously combined successfully with wheat exome capture (Gardiner *et al.*, 2016).

This assay brings re-sequencing of the entirety of the high confidence gene-associated portion of wheat within the reach of the wheat community. Our multiplexing analysis defined more than 1.8 million positions across eight diverse samples, where each of the samples had a minimum of 5X coverage to allow comparison, and variation was observed between samples. This level of SNP information will allow refinement of key genetic regions linked to traits and enable researchers to pinpoint phenotype-inducing SNPs more precisely. Finally, we predict that our optimization of the protocol for this large-scale capture using an island approach will allow us to sequence >90% of the gene-space of up to four wheat accessions on a single HiSeq4000 lane and twenty accessions on a NovaSeq S1 flow cell to a minimum of 5X (>80% at >10X). Our capture probe set design is publicly available and can also be ordered directly from NimbleGen via the Roche website (http://sequencing.roche.com/en/products-solutions/by-category/target-enrichment/shareddesigns.html).

### Potential Implications

We have previously demonstrated the use of sequence capture to allow the study of both genotype and DNA methylation across targeted regions in wheat (Gardiner *et al.*, 2015). Using bisulfite treatment after sequence capture, DNA methylation analyses can be performed using the same probe sets that are implemented for genotyping (Olohan *et al.*, 2018). Moreover, we have demonstrated the use of sequence capture that was designed using the reference wheat variety Chinese Spring to analyze diverse landraces from the Watkins collection (Gardiner *et al.*, 2018) and even highly divergent ancient wheat diploid progenitors with high efficiency (Gardiner *et al.*, 2014; Grewal *et al.*, 2017). As such, it is likely that the capture probe sets defined here could not only effectively enable re-sequencing of the high confidence genes of bread wheat lines, they could be used to further epigenetics research and research across a broader variety of wheat accessions than we tested here. The integration of more diverse wheat diploid and tetraploid progenitor material into the design will also allow broad applicability of the probe sets to varieties beyond bread wheat and also to synthetic wheat lines, constructed from diploid and tetraploid progenitors, that are becoming increasingly popular in the wheat community.

## Methods

### Developing the capture probe design space from its target regions (Figure 1)

Initially the target gene/promoter sequences from each wheat reference genome (TGAC-Chinese Spring, IWGSC-Chinese Spring, *Ae. tauschii* and Emmer) were processed independently of one another. For each gene/promoter set (Figure 1) gene/promoter sequences were aligned to themselves using BLASTN with a maximum e-value of 1e-5, minimum sequence identity of 95% and minimum match length of 100bp. Here, non-redundant sequences with no BLASTN alignments were taken forward directly (known as NR-sequences). Any full or partial sequences that aligned to other sequences in the gene/promoter set were extracted and BLASTclust was used to cluster these redundant sequences by similarity allowing the longest representative sequence per alignment group to be identified and combined with the NR-sequences to be taken forward. Furthermore, if parts of otherwise NR-sequences were redundant and removed but were then outputted from the BLASTclust alignment as a representative non-redundant sequence, these fragments were then re-integrated back into their sequence of origin. This generated a complete, re-assembled where possible, set of NR-sequences.

The complete set of NR-sequences was aligned to the wheat chloroplast/mitochondria genomes using BLASTN, with the same parameters used previously, and regions or sequences showing hits were removed. Dustmasker was then implemented to annotate and low-complexity regions as lower case; later during probe design, probes with low-complexity regions of 40 bp or more were disregarded. Finally, NR-sequences less than 120 bp in length were removed from the sequence set. This yielded individual probe set design spaces for TGAC-wheat, IWGSC-wheat, Emmer and *Ae. Tauschii.*

These sequence sets were then compared to identify species overlap using a BLASTN alignment with the same parameters used previously. Emmer and *Ae. Tauschii* design spaces were compared to the TGAC wheat design space and to each other and any unique Emmer/*Ae. Tauschii* sequences were combined with the TGAC wheat design space. The Chinese Spring IWGSC design space was then compared to this TGAC/Emmer/Tauschii design space and unique sequences were combined. Finally, fragments of less than 75 bp in length were removed to generate a TGAC/Emmer/Tauschii/IWGSC gene and promoter design space.

### Exonic mature miRNA selection

pre-miRNA sequences that were annotated using IWGSC Refseq1.v1 were mapped using BLA on the chromosome sequences of the same genome in order to obtain their exact genomic locations. The pre-miRNA start and end alignment positions were compared with the boundaries of the exons in the annotated set of genes from IWGSC and TGAC using an in-house python script. After classification, mature miRNAs that were identified from pre-miRNAs whose start and end sites were both located in the same exonic region were directly called as exonic miRNAs. In the case of pre-miRNAs with start and end sites located on different regions, mature miRNA locations were taken into account by comparing their start and end sites with exons. If the whole mature miRNA sequence was located within an exon, they were also called as exonic miRNAs. These sequences were added to the promoter capture design space.

### Characteristics of the exome capture kit

The final gene and promoter capture design space (785,914,746 bp) was processed by NimbleGen for probe design. The NimbleGen probe set manufacturing platform has a maximum capacity of 2.16 million probes that are typically 50-100 nucleotides in length with an average of 75 bp i.e. maximum actual probe space ~162 Mb. Typically probes overlap one another to most optimally cover the target design space, however from previous analyses we observed that a single 120 bp probe can enrich up to 500 bp routinely with adequate sequencing coverage (Gardiner *et al.*, 2015). As such, we requested that probes be tiled across our design space using an “island strategy” where probes are spaced at intervals, to most evenly cover the design-space. This resulted in probes being tiled across our design space at an average spacing of 120 bp from the 5’ start of a probe to the 5’ start of the next probe. The best probe within a 20 bp window of this start location was selected to minimize low complexity sequence in probes.

### Sample library preparation and in solution captures

Genomic DNA was extracted from Chinese Spring and the eight CIMMYT lines (21-day seedling leaf tissue) using the Qiagen DNeasy plant mini kit. For Chinese Spring, 1μg aliquots of the genomic DNA, each in a total volume of 55μl, were sheared for 2×60s using a Covaris S2 focused-ultrasonicator (duty cycle 10%, intensity 5 and 200 cycles per burst using frequency sweeping). For the eight CIMMYT samples, 1 μg of each genomic DNA sample, in a total volume of 55 μl, was sheared for 1×60s using a Covaris S2 focused-ultrasonicator (duty cycle 5%, intensity 5 and 200 cycles per burst using frequency sweeping). The fragmented DNA was directly used as input for library preparation. The NimbleGen SeqCap EZ Library SR User’s Guide (Version 5.1, September 2015) was followed for all steps with the modifications listed below.

The dual size selection of the pre-capture libraries was adjusted to account for the larger shearing sizes. For Chinese Spring the volumes were 45 μl and 20 μl for right and left size selection, respectively. For the CIMMYT samples the volumes were 40 μl and 20 μl. Five cycles of amplification were used for the pre-capture PCR. The capture input for the Chinese Spring captures was 2 μg DNA and 1.4 μg for the CIMMYT captures. A higher input was used for Chinese Spring to increase final library yield, but it was subsequently found that 1.4μg was sufficient. Since the input DNA was derived from wheat, 1μl of Developer Reagent Plant Capture Enhancer (NimbleGen) was added per 100 ng input in the hybridisation step instead of COT human DNA. The SeqCap HE Universal Oligo (NimbleGen) and SeqCap HE Index Oligo pool (NimbleGen) were added separately and the volume of SeqCap HE Universal Oligo was adjusted to 3.4 μl and 2.8 μl for the Chinese Spring and CIMMYT captures, respectively. This increase in volume was to account for the higher DNA inputs. Finally, for the final post-capture PCR, 14 cycles were used for the Chinese Spring captures and 12 cycles for the CIMMYT captures. The cycle number was reduced to 12 cycles as this still produced a high enough yield sequencing.

### Quality control for the promoter and gene capture

An initial assessment of library yield was made using Qubit High Senstivity double stranded DNA assays (Invitrogen). Fragment size distribution was determined from Bioanalyser High Sensitivity DNA (Agilent) data. Prior to sequencing the libraries were quantified by qPCR, using an Illumina Library Quantification Kit (KAPA) on an Applied Biosystems StepOne system.

To assist in the determination of enrichment efficiency post-capture, we designed qPCR primers that cover probe targets. These are as follows for the gene capture; forward “CCGAGCCTCATAGTCAGGAG” and reverse “TGGGAAAACTGATCCCAGTC”. For the promoter capture probe set the recommended primers are as follows; forward “CTGTTTGTTTTGAGCGCGTC” and reverse “TGGCTTCGCGAAACTGAAAA”. The polymerase master mix from the Illumina Quantification Kit and StepOne system were used to perform the enrichment qPCR. The qPCR reaction conditions were as follows, 95 °C 10 minutes and forty cycles of 95 °C for 10 seconds, 72 °C for 30 seconds, and 60 °C for 30 seconds. The qPCR was performed on aliquots of the capture library pre and post-capture; after first diluting the aliquots to the same ng/μl concentration. The ΔCT between the pre- and post-capture of successful gene capture ranged from 4 to 5. For promoter captures the ΔCT ranged from 3 to 4.

### Illumina DNA sequencing of gene and promoter captures

For the Chinese Spring sample, four technical replicate barcoded libraries were pooled for the gene capture and a further four were pooled for the promoter capture. The final two capture libraries were pooled using a ratio of 33%:66% promoter-to-gene to reflect the different size targets of the probe sets. This pool was then sequenced on a single HiSeq4000 lane. This generated 2×150bp reads. For the eight CIMMYT lines, the same barcoded libraries were used for individual gene and promoter captures, therefore these captures were sequenced separately across multiple HiSeq4000 lanes. The read data produced was equivalent to 1½ and 2½ HiSeq4000 lanes for the promoter and gene capture, respectively.

Separate sequence capture experiments were conducted at KSU Integrated Genomics Facility using the promoter-2 capture assays following the same capture protocol with the following modifications. The capture reaction was performed on a set of 22 pooled samples barcoded using dual indexes. These samples were pooled into a larger pool of 96 barcoded sequence capture libraries and sequenced using 2 × 150 bp sequencing run on the S1 flow-cell of NovaSeq 6000 system.

### Optimizing the capture protocol

Here the standard capture protocol described above is followed, but with the following modifications: average insert size was <300bp rather than 350 or 500 bp for the Chinese Spring and CIMMYT captures, respectively; the KAPA Hyper library preparation kit rather than the KAPA HTP preparation kit was used; adapter ligation was performed overnight at 4 °C rather than for 15 minutes at 20 °C; and an additional wash of the bead bound-capture library with 50 μl PCR-grade water was included. This was part of a development of a methylation capture protocol.

### Initial sequence data analysis

Mapping analyses of sequencing reads were carried out using BWAmem (version 0.7.10) (Li and Durbin, 2009) and HISAT2 (Kim *et al.*, 2015). Paired-end reads were mapped and only unique best mapping hits were taken forward. Mapping results were processed using SAMtools (Li *et al.*, 2009) and any non-uniquely mapping reads, unmapped reads, poor quality reads (< Q10) and duplicate reads were removed. SNP calling was carried out using the GATK Unified genotyper (after Indel realignment), which was used with a minimum quality of 50 and filtered using standard GATK recommended parameters, a minimum coverage of 5X and only homozygous SNPs were selected as defined by GATK i.e. allele frequency in >80% of the sequencing reads (McKenna *et al.*, 2010). Furthermore, if three or more SNPs occurred within a 10 bp window these were filtered out from the calls.

## Availability of supporting data and materials

The sequencing data sets supporting the results of this article are available in the European Nucleotide Archive repository, study PRJEB27620. The final design space for the capture probes sets and the locations of the capture probes on this design space are available from the Grassroots Data Repository (http://opendata.earlham.ac.uk/wheat/under_license/toronto/Gardiner_2018-07-04_Wheat-gene-promoter-capture/). The target locations of the capture probe sets on the Chinese Spring IWGSC RefSeqv1 i.e. the high confidence gene and promoter sequences, are detailed in supporting files 2, 3 and 4.

### Supporting data

File 1: Supplementary_data.docx

Includes Supplementary Figures S1-S4, Supplementary Tables S1-S4

File 2: Gene-capture-HC-targets.bed

File 3: Prom-capture-HC-targets.bed

File 4: Prom-capture-HC+5UTR-targets.bed

## Declarations

### List of abbreviations

CIMMYT: International Maize and Wheat Improvement Center (Centro Internacional de Mejoramiento de Maíz y Trigo)
IWGSC: International Wheat Genome Sequencing Consortium
miRNA: Micro RNA
NR: Non-redundant
PCR: Polymerase Chain Reaction
qPCR: Quantitative Polymerase Chain Reaction
SNP: Single Nucleotide Polymorphism
TGAC: The Genome Analysis Centre (now known as the Earlham Institute)
TSS: Transcription Start Site
UTR: Untranslated Region

## Consent for publication

All plants used in this study were grown in controlled growth chambers complying with Norwich Research Park guidelines.

## Authors’ contributions

The capture probe set design, Chinese Spring and CIMMYT line bioinformatic validation and manuscript preparation was performed by LG. The project was designed, planned and conducted by LG and AH. Plant growth, DNA extractions, library prep and sequence capture were performed by TB with assistance from LC. TR assisted with capture probe design. The CIMMYT material was contributed by S.S. E.A. and H.B. contributed to the promoter capture assay design. A.A. conducted sequence capture and NGS sequencing, K.J. contributed to analysing promoter capture data for the 22-plex test. All authors read and approved the final manuscript.

## Acknowledgements

We thank Genomic Pipelines at the Earlham Institute for DNA sequence generation and Cristobal Uauy for his advice regarding capture design. We would also like to thank Xingdong Bian, Simon Tyrrell and Robert Davey for their help adding our sequence data onto the Grassroots Data Repository.

## References

Avni R, Nave M, Barad O, Baruch K, Twardziok SO, Gundlach H, Hale I, Mascher M, Spannagl M, Wiebe K et al. 2017. Wild emmer genome architecture and diversity elucidate wheat evolution and domestication. Science 357(6346): 93–97

Baird NA, Etter PD, Atwood TS, Currey MC, Shiver AL, Lewis ZA, Selker EU, Cresko WA and Johnson EA. 2008. Rapid SNP discovery and genetic mapping using sequenced RAD markers, PLoS ONE 3:e3376

Brenchley R, Spannagl M, Pfeifer M, Barker GLA, D’Amore R, Allen AM, McKenzie N, Kramer M, Kerhornou A, Bolser D et al. 2012. Analysis of the bread wheat genome using whole-genome shotgun sequencing. Nature 491, 705–710

Cavanagh CR, Chao S, Wang S, Huang BE, Stephen S, Kiani S, Forrest K, Saintenac C, Brown-Guedira GLB, Akhunova A et al. 2013 Genome-wide comparative diversity uncovers multiple targets of selection for improvement in hexaploid wheat landraces and cultivars. PNAS, 110(20): 8057–8062

Clavijo BJ, Venturini L, Schudoma C, Accinelli GG, Kaithakottil G, Wright J, Borril P, Kettleborough G, Heavens D, Chapman H et al. 2017. An improved assembly and annotation of the allohexaploid wheat genome identifies complete families of agronomic genes and provides genomic evidence for chromosomal translocations, Genome Research 27(5):885–896

De Wit P, Paspeni MH and Palubi SR. 2015. SNP genotyping and population genomics from expressed sequences – current advances and future possibilities, Mol Ecol, 24:2310–2323.

Dubcovsky J and Dvorak J. 2007. Genome plasticity a key factor in the success of polyploid wheat under domestication. Science 316, 1862–1866

Dvorak J, Akhunov ED, Akhunov AR, Deal KR and Luo MC. 2006. Molecular characterization of a diagnostic DNA marker for domesticated tetraploid wheat provides evidence for gene flow from wild tetraploid wheat to hexaploid wheat, Mol. Biol. Evol. 23, 1386–1396

Gardiner, LJ., Gawronski, P., Olohan, L., Schnurbusch, T., Hall, N. and Hall, A., 2014. Using genic sequence capture in combination with a syntenic pseudo genome to map a deletion mutant in a wheat species. The Plant Journal, 80;5, 895–904.

Gardiner LJ, Quinton-Tulloch M, Olohan L, Price J, Hall N and Hall A. 2015. A genome-wide survey of DNA methylation in hexaploid wheat. Genome Biology 16: 273

Gardiner LJ, Bansept-Basler P, Olohan L, Joynson R, Brenchley R, Hall N, O’Sullivan DM and Hall A. 2016. Mapping-by-sequencing in complex polyploid genomes using genic sequence capture: a case study to map yellow rust resistance in hexaploid wheat. The Plant Journal 87 (4), 403–419

Gardiner LJ, Joynson R, Omony J, Rusholme-Pilcher R, Olohan L, Lang D, Bai C, Hawkesford M, Salt D, Spannagl M et al. 2018. Hidden variation in polyploid wheat drives local adaptation. bioRxiv: https://doi.org/10.1101/217828

Grewal S, Gardiner L, Ndreca B, Knight E, Moore G, King IP and King J. 2017. Comparative Mapping and Targeted-Capture Sequencing of the Gametocidal Loci in Aegilops sharonensis. Plant Genome 10. doi:10.3835/plantgenome2016.09.0090

Jordan K, Wang S, Lun Y, Gardiner L, MacLachlan R, Hucl P, Wiebe K, Wong D, Forrest K, IWGSC et al. 2015. A haplotype map of allohexaploid wheat reveals distinct patterns of selection on homoeologous genomes. Genome Biol. 16, 48.

Kim D, Langmead B and Salzberg SL. 2015 HISAT: a fast spliced aligner with low memory requirements. Nat. Methods, 12, 357–60.

Krasileva K, Vasquez-Gross HA, Howell T, Bailey P, Paraiso F, Clissold L, Simmonds J, Ramirez-Gonzalez RH, Wang X, Borril P et al. 2017. Uncovering hidden variation in polyploid wheat. PNAS, 114 (6) E913–E921; doi:10.1073/pnas.1619268114

Luo MC, Gu YQ, Puiu D, Wang H, Twardziok SO, Deal KR, Huo N, Zhu T, Wang L, Wang Y, McGuire PE et al. 2017. Genome sequence of the progenitor of the wheat D genome *Aegilops tauschii*, Nature 551: 498–502

Li H and Durbin R. 2009. Fast and accurate short read alignment with Burrows-Wheeler transform. Bioinformatics 25, 1754–1760

Li H, Handsaker B, Wysoker A, Fennell T, Ruan J, Homer N, Marth G, Abecasis G, Durbin R and 1000 Genome Project Data Processing Subgroup. 2009. The Sequence Alignment/Map format and SAMtools. Bioinformatics 25, 2078–2079

Li X, Zhu J, Hu F, Ge S, Ye M, Xiang H, Zhang G, Zheng X, Zhang H, Zhang S et al. 2012. Single-base resolution maps of cultivated and wild rice methylomes and regulatory roles of DNA methylation in plant gene expression. BMC Genomics 13, 300

McKenna A, Hanna M, Banks E, Sivachenko A, Cibulskis K, Kernytsky A, Garimella K, Altshuler D, Gabriel S, Daly M and DePristo MA. 2010. The Genome Analysis Toolkit: A MapReduce framework for analyzing next-generation DNA sequencing data. Genome Res 20, 1297–1303

Wicker T, Gundlach H, Spannagl M, Uauy C, Borrill P, Ramirez-Gonzalez R, De Oliveira R, IWGSC, Mayer K, Paux E and Choulet F. 2018. Impact of transposable elements on genome structure and evolution in bread wheat. In press

Winfield MO, Allen AM, Wilkinson PA, Burridge AJ, Barker GLA, Coghill J, Waterfall C, Wingen LU, Griffiths S and Edwards KJ. 2017. High density genotyping of the A.E. Watkins Collection of hexaploid landraces identifies a large molecular diversity compared to elite bread wheat, Plant Biotechnology Journal, 16(1): 165–175

